# Healthy human eyes’ misaligned optical components: Binocular Listing’s law

**DOI:** 10.1101/2025.06.23.661014

**Authors:** Jacek Turski

**Author notes:** Jacek Turski contributed all work to this manuscript.

## Abstract

The healthy human eye’s optical components are misaligned. Although important in studying vision quality, it has been overlooked in research on binocular and oculomotor vision. This study presents the construction of ocular torsion in the binocular system that incorporates the fovea displaced from the posterior pole and the lens tilted away from the eye’s optical axis. When the eyes’ binocular posture changes, each eye’s torsional position transformations, computed in the framework of Rodrigues’ vector, are visualized in *GeoGebra* simulations. Listing’s law, important in oculomotor control by constraining a single eye redundant torsional degree of freedom, is ab initio formulated for bifoveal fixations in the binocular system with misaligned optical components for the fixed upright head. It leads to the configuration space of binocularly constrained eyes’ fixations, including the noncommutativity rule. This formulation modifies the Listing plane of the straight-ahead eye’s primary position by replacing it with the binocular eyes’ posture corresponding to the empirical horopter’s abathic distance fixation, a unique bifoveal fixation for which the longitudinal horopter is a straight frontal line.

Notably, it corresponds to the eye muscles’ natural tonus resting position, which serves as a zero-reference level for convergence effort. Supported by ophthalmology studies, it revises the elusive neurophysiological significance of the Listing plane. Furthermore, the binocular constraints couple 3D changes in the eyes’ orientation and, hence, torsional positions during simulations with *GeoGebra*’s dynamic geometry. The binocular Listing’s law developed here can support this coupling, which is important in oculomotor control. The results obtained in this study should be a part of the answers to the questions posted in the literature on the relevance of Listing’s law to clinical practices.

**Author summary:** Our eye optical components are misaligned: the fovea is displaced from the eye’s posterior pole, and the lens is tilted away from the optical axis. Listing’s law, important in oculomotor control, has not only overlooked the misaligned eye’s optics but was also formulated for a single eye, with a later ad hoc extension added for binocular vision. The purpose of Listing’s law is to constrain the eye’s redundant torsional degrees of freedom, thereby supporting neural processing in the development of our spatial understanding by controlling the noncommutativity of the eye’s rotations. This goal cannot be fully met because Listing’s law is monocular, but we acquire an understanding of the scene through bifoveal fixations on objects. In this work, I construct ocular torsion that accounts for the eye’s misaligned optics and incorporate it into Listing’s law. It directly leads to its first ab initio consistent binocular formulation, which is visualized in a computer simulation. Supported by ophthalmological studies, it revises the still elusive neurophysiological significance of the Listing plane, the basic ingredient of Listing’s law. It also resolves the persistent lack of a generally accepted explanation for Listing’s law. The results of this study are likely to be important in the ongoing discussion in the literature regarding the relevance of Listing’s law to clinical practices.

## Introduction

In healthy human eyes, the fovea is displaced on the retina from the posterior pole, and the crystalline lens is tilted away from the eye’s optical axis. This asymmetry has been studied in many clinical publications, usually decomposed along temporal-nasal and inferior-superior axes or horizontal and vertical segments [1–5]. The fovea’s anatomical displacement, relatively stable in the human population [6], and the cornea’s asphericity contribute to optical aberrations, and the lens tilt partially compensates for these aberrations [7–10]. Nevertheless, it has been overlooked in binocular and oculomotor vision research.

The binocular system with the 2D asymmetric eye (AE) model, which comprises fovea and lens horizontal misalignments, has been developed in [11]. Then, in [12, 13], its consequences for stereopsis and visual space geometry were elucidated. Based on these studies, a binocular system with a 3D AE model was formulated in [14], incorporating the fovea and lens’ horizontal and vertical misalignment angles. The consequences and functional role of vertical misalignment of the eye’s optics included an explanation of the observed inclination of the vertical horopter, as well as the proposition of an ab initio binocular formulation of Listing’s law.

The rationale behind this article is to focus solely on Listing’s law. This will enable the definitive ab initio binocular modification of Listing’s law within the framework of the 3D AE model, supported by ophthalmology and experimental studies, as well as new geometric formulations and numerical simulations. The unique results include, for a stationary upright head, clarifying the configuration space of binocularly constrained eyes’ fixations, including the noncommutativity rule.

Listing’s law states that for a stationary, upright head, from a reference position, the single eye rotation axes lie in a plane referred to as the displacement plane. The displacement planes associated with different reference positions do not coincide. When the reference position is perpendicular to the displacement plane, the plane is referred to as the Listing plane, and the unique reference position is the primary position. Further, for any eye’s tertiary position, the displacement plane is obtained by rotating the Listing plane by the half-angle of gaze eccentricity. This corollary of Listing’s law is known as the half-angle rule. Listing’s law holds for the eyes in optical infinity.

Improvements in eye movement recording techniques have shown that during convergence on a near object, the eyes’ rotation axes remain confined to planes that are rotated temporally by angles proportional to the vergence for the right and left eye [15–19]. This binocular extension is known as L2. However, from the experimental data in those references, the coefficient of proportionality ranges from 0.2 to 0.5. Thus, the L2 law is an ad hoc extension of Listing’s law.

It follows from the above discussion that, apart from the overlooked misaligned optical components of healthy human eyes, the most striking shortcoming of Listing’s law is the lack of a consistent binocular formulation. This is further evident in the commonly expressed purpose that Listing’s law constrains a single eye’s redundant torsional degrees of freedom to support neural processing in building up our spatial understanding [20]. This goal cannot be fully met because Listing’s law is monocular, but we acquire the scene’s understanding with bifoveal fixations on objects. Further, the 3D binocular movements of the eyes couple their torsional positions [21].

Not surprisingly, there are many controversies regarding Listing’s law’s nature and functions, as discussed in [21–23], for example. In particular, there is no generally accepted explanation for Listing’s law [21], which is still true today.

The AE comprises the nodal point and the image plane passing through the eye’s rotation center. The image plane orientation represents the lens’ tilt, and the retina is replaced by its projection through the nodal point into the image plane. The AE incorporates the eye’s anatomy and physiology in supporting biologically mediated aspects of binocular vision that it intends to model. Thus, it satisfies the law of parsimony: the best functional eye models include only necessary attributes for accomplishing the models’ purpose [24].

The AE model has a curious predecessor in the field of vision science. In 1959, Frederick Verhoeff, a distinguished ophthalmologist, wrote in [25]: “[…] because we see straight lines with spherical retinas, mathematically, the eyes can be replaced with ‘miniature pinhole cameras’.” He placed two cameras at the interocular distance, pointing straight forward, so their screens are in the same plane. Further, Verhoeff accounted for the alpha angle—the angular displacement of the fovea on the retina from the posterior pole, which he believed might benefit binocular vision. To do this, he first assumed that the distance and direction of the corresponding points from the foveae (the optical centers) were the same on both camera screens. Then, he moved the principal corresponding points, i.e., the foveae, on the screen of the right camera to the right and on the screen of the left camera to the left so that the camera would have an equal non-zero angle alpha. Of course, his eye model did not include a tilted lens (angle beta), which contributes to the eye’s aplanatic design by canceling out some of the aberrations caused by the alpha angle, cf. Fig. 1 (b).

**Fig 1.**
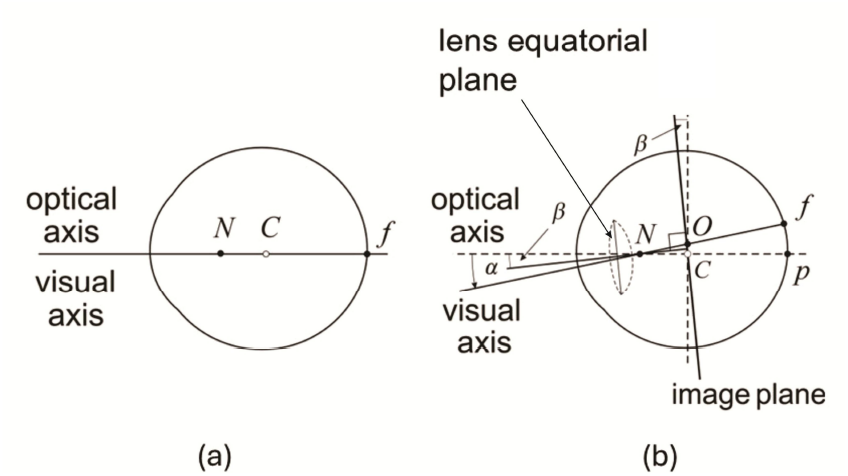
The AE model. (a) The reduced eye. (b) The right AE. *N* is the nodal point, *C* is the eye’s rotation center, *f* is the fovea, and *p* is the posterior pole. The angles *α* and *β* model horizontal misalignment of the fovea and the lens. The image plane is parallel to the lens’s equatorial plane and passes through the eye’s rotation center. *O* is the projection of the fovea into the image plane—the optical center on the image plane.

Interestingly, three years after Verhoeff introduced his eye model of the flat-screen pinhole camera with angle alpha, he used it in [26] to explain Kundt and Tschermak-Seysenegg’s illusions.

In the remaining part of this section, I explain the AE model that incorporates the healthy human eye’s misaligned fovea and lens and review the main results obtained in [11–14]. It provides background to help the reader understand the rest of the paper.

However, before I first explain the AE model, I will recall the notion of stereopsis. If for two retinal elements, one in each eye, a localized stimulus is perceived in a single direction irrespective of whether it reaches only one element or the other or both simultaneously, they are considered to be corresponding elements of zero disparity determined by the nonius paradigm [27]. The locus of points in the binocular field, such that each point projects to a pair of corresponding retinal elements, is known as the horopter. The horizontal correspondence between disparate 2D retinal images of our laterally separated two eyes organizes visual perception of objects’ form and their location in 3D space, i.e., stereopsis and spatial relations [28, 29].

### 2D AE model

The 2D AE model shown in Fig. 1 (b), which extends the standard axially symmetric single refractive surface schematic eye model (reduced eye) in Fig. 1 (a), was introduced in [11] for horizontally misaligned fovea and lens. Because the eyes are horizontally displaced in the head, horizontal misalignment impacts stereopsis.

The lens in the AE is represented by (i) the *nodal point* located on the optical axis 0.6 cm anterior to the eye’s rotation center and (ii) the *image plane* parallel to the lens’s equatorial plane and passing through the eye’s rotation center. It provides a minimal requirement to account for the lens tilt. The fovea in AE is displaced horizontally from the posterior pole in the temporal direction by *α* degrees, and the lens is horizontally tilted away from the eye’s optical axis by *β* degrees. Both angles are measured at the nodal point. The typical values of the angles in the human population are *α* = 5.2° and *β* = 3.3°.

The impact of misalignment on the longitudinal horopter shape was studied in [12].

Fig. 2 (a) shows simulations of typical shapes of the horopters; an ellipse for near fixation and a hyperbola for farther fixation, divided by the straight frontal line horopter for the symmetric fixation *F*_*a*_ with coplanar lens equatorial planes parallel to the coplanar image planes, as explained in Fig. 2 (b). The simulated horopters in Fig. 2 (a) resemble empirical horopters first measured by Hillebrand about 130 years ago, but not using the nonius paradigm.

**Fig 2.**
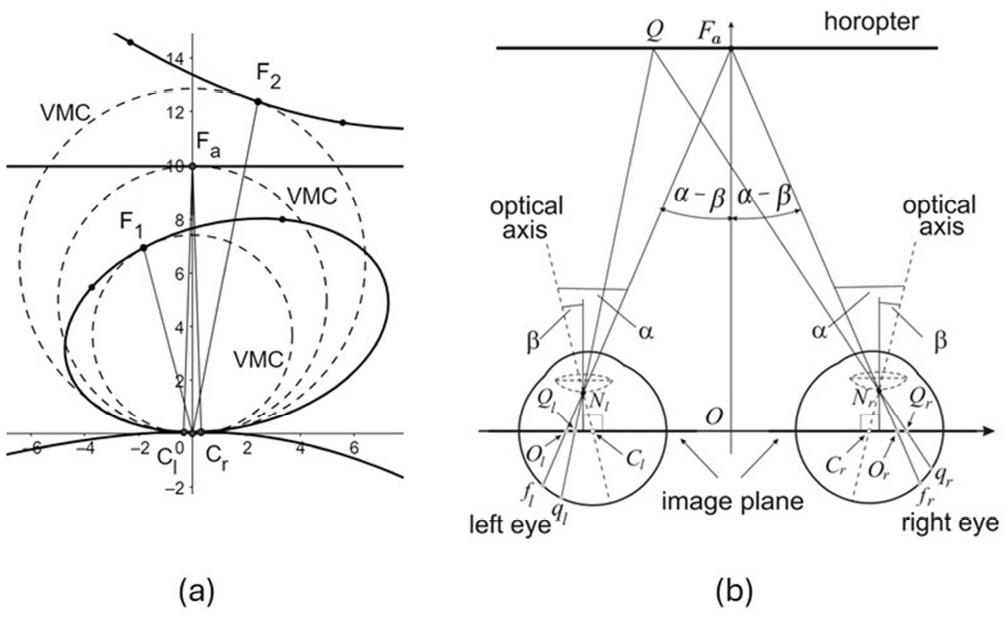
Horopters in the binocular system with AEs. (a) Horopteric conics rendered in *GeoGebra* are shown in continuous lines for the anthropomorphic dimensions of the binocular system with the AEs for the typical values of *α* = 5.2° and *β* = 3.3° in the human population. The Vieth-Müller circles (VMC) shown in dashed lines are geometric horopters for the reduced eye. (b) The fixation *F*_*a*_ corresponds to the AEs’s posture in which the coplanar image planes are parallel to the lens coplanar equatorial planes. The horopter through *F*_*a*_ is a straight frontal line. The fixation *F*_*a*_ projects to the foveae *f*_*r*_ and *f*_*l*_ in the retina and the optical centers *O*_*r*_ and *O*_*l*_ in the image plane. The point *Q* on the horopter projects to the corresponding retinal points *q*_*r*_ and *q*_*l*_, and to points *Q*_*r*_ and *Q*_*l*_ in the image plane. Although |*f*_*l*_*q*_*l*_| ≠ |*f*_*r*_*q*_*r*_| on the retina, |*O*_*l*_*Q*_*l*_| = |*O*_*r*_*Q*_*r*_| on the image plane.

For angles *α* and *β*, the distance to *F*_*a*_ is derived in [12] as follows:

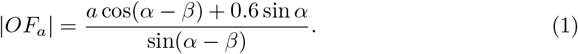

Here, 2*a* = 6.5 cm is the ocular separation, 0.6 cm is the distance of the nodal point from the eyeball’s rotation center, and *O* is the head coordinate system origin. With *a* = 3.25 cm, *α* = 5.2° and *β* = 3.3°, |*OF*_*a*_| is about 99.6 cm, which agrees with the average abathic distance fixation of about 1 meter.

It was demonstrated in [13] that for fixation *F*_*a*_, iso-disparity curves that provide retinal spatial coordinates, where the horopter is the zero-disparity curve, are straight frontal lines. This unique binocular position is referred to as the eyes’ resting posture (ERP). For all other fixations in the horizontal visual plane, the iso-disparity curves are ellipses or hyperbolas.

The main original contributions to binocular vision resulting from the studies mentioned above are as follows:

1. The asymmetric retinal correspondence—the corresponding elements are compressed in the temporal retinae relative to those in the nasal retinae—which is person-dependent and difficult to measure [30], is represented under projection through the nodal points by a symmetric distribution of points in the image plane of the ERP. This representation depends on the AE’s horizontal misalignment angles. The correspondence symmetric distribution on the image plane is preserved for all binocular fixations [12].
2. The asymmetry of the retinal correspondence is caused by the horizontal misalignment of the fovea and the lens in healthy human eyes. It is demonstrated in [13] by the spatial iso-disparity distribution being *universal* for the AE’s angles of horizontal misalignment. It means the iso-disparity line distribution for a fixed *α* is invariant to *β* values in the AE.

Furthermore, in [13], the framework of Riemannian geometry examined the global aspects of phenomenal spatial relations, including visual space variable curvature and a finite horizon. The global aspects of stereopsis are crucial in perception, as exemplified by the coarse disparity contribution to our impression of being immersed in the ambient environment despite receiving 2D retinal projections of the light beams reflected by spatial objects [31, 32].

### 3D AE model

The 2D AE is extended in [14] by including vertical misalignment based on the clinical studies discussed before. The typical in the human population asymmetry angles for the fovea’s horizontal and vertical displacements from the posterior pole are *α* = 5.2° and *γ* = −2° and the lens’ horizontal and vertical tilts relative to the optical axis are *β* = 3.3° and *ε* = −1°, explained in Fig. 3 for the right AE. The orientation of the axes determines the signs of angles.

**Fig 3.**
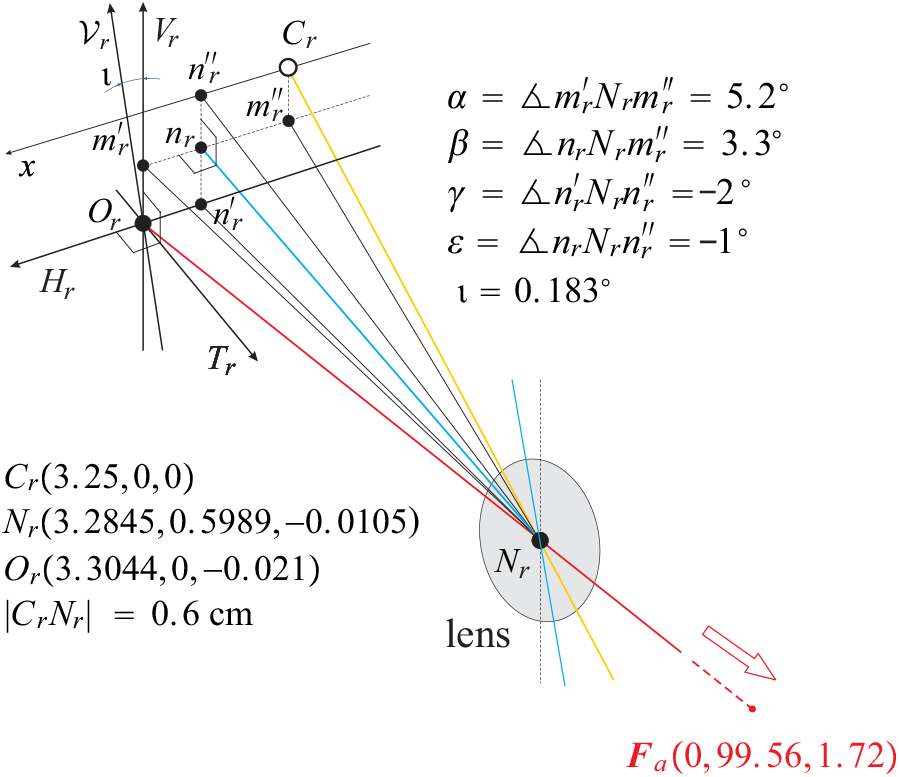
Horizontal and vertical angles of misalignment in the right AE. The right 3D AE view is shown near the optical center *O*_*r*_ and the eye’s rotation center *C*_*r*_. *N*_*r*_ is the nodal point, the yellow line is the eye’s optical axis, the red line is the visual axis fixating *F*_*a*_, and the blue line through *n*_*r*_ and *N*_*r*_ is the lens’s optical axis. The AE’s image plane coordinates consist of the horizontal axis *H*_*r*_ and the vertical axis *V*_*r*_, both of which are attached at *O*_*r*_, the projected fovea in the image plane, cf. Fig. 1 (b). They are complemented by the perpendicular axis *T*_*r*_ parallel to the lens’s optical axis. *H*_*r*_ and *V*_*r*_ are the retinal horizontal and vertical correspondence axes, cf. Fig. 4.

Similarly to 2D AE, the binocular system with AEs has a unique fixation for the given asymmetry angles *α, β, γ*, and *ε*, such that the resulting posture consists of vertical coplanar image planes parallel to the coplanar lenses’ equatorial planes. For the values of asymmetry angles listed above, the fixation of the special posture is *F*_*a*_(0, 99.56, 1.72), with coordinates in centimeters, corresponding to an average abathic distance fixation of approximately 1 meter; this is the ERP.

In Fig. 3, the axis *H*_*r*_ is the image plane axis of retinal horizontal correspondence for the right AE–the location of the corresponding points that are used to construct the longitudinal iso-disparity curves in Fig. 4 and simulated in Fig. 6. In contrast, the retinal vertical correspondence is not defined on the vertical axis *V*_*r*_ in Fig. 3 but instead on the axis denoted by *V*_*r*_ that is tilted from the true vertical axis *V*_*r*_ by *ι*= 0.183° with its top in the temporal direction; see also Fig. 4.

**Fig 4.**
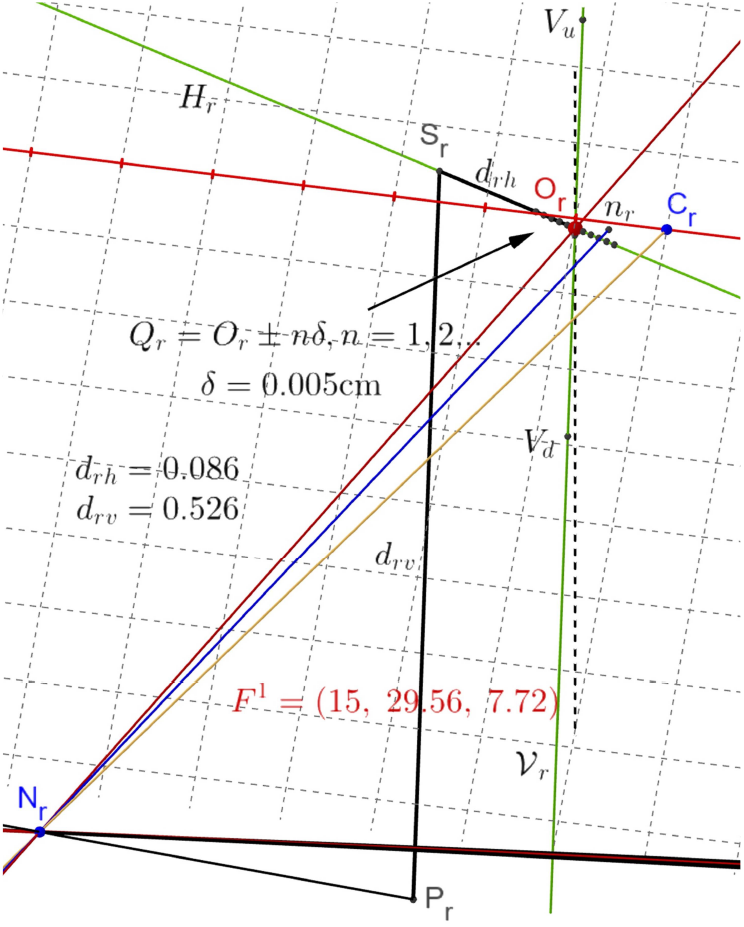
The construction of the iso-disparity conics and vertical horopter. The construction is explained for the right eye fixating on *F* ^1^(15, 29, 56, 7.72). The visual axis is the red line passing through the nodal point *N*_*r*_ and the optical center *O*_*r*_. The optical axis is the yellow line from *C*_*r*_ and through *N*_*r*_. The blue line through *N*_*r*_ and *n*_*r*_ is the lens’s optical axis, perpendicular to the image plane. The green line *H*_*r*_ through *O*_*r*_ is the foveal horizon, with the horizontal disparity points placed 0.005 cm apart. The green line *V*_*r*_ through *O*_*r*_ is the axis of vertical retinal correspondence tilted from the true vertical (dashed line) by 0.183°. *P*_*r*_ is the projection through *N*_*r*_ of the point *P* in the binocular field into the right image plane. *d*_*rh*_ is the horizontal disparity and *d*_*rv*_ is the vertical disparity along the corresponding axes.

**Fig 5.**
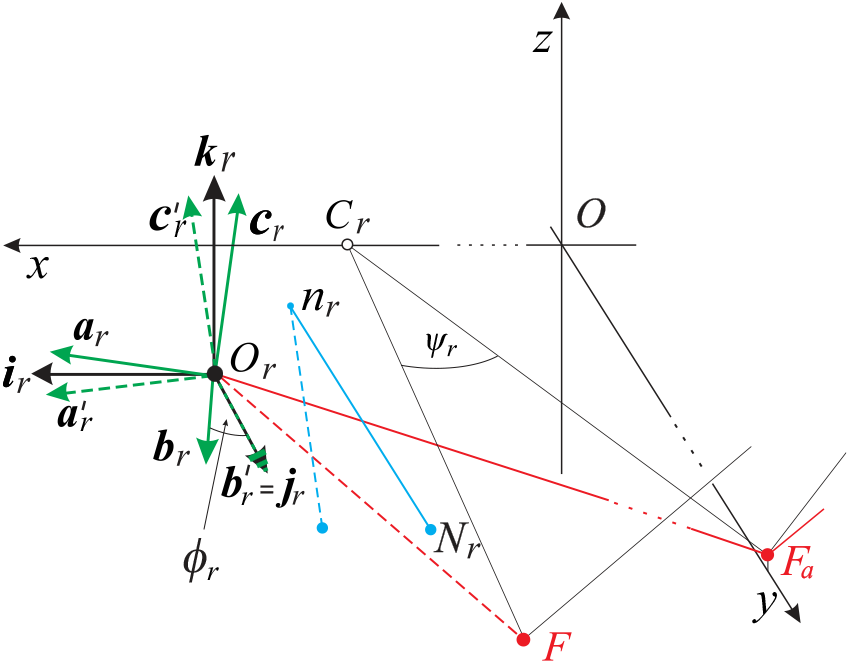
The definition of ocular torsion in the right AE. The directions of black frame (***i***_*r*_, ***j***_*r*_, ***k***_*r*_) vectors at *O*_*r*_, agrees with the direction of lines *H*_*r*_, *T*_*r*_ and *V*_*r*_ in Fig. 3 and with the directions of head coordinates; *x, y, z*. The green frame (***a***_*r*_, ***b***_*r*_, ***c***_*r*_) is the rotated black frame when the fixation axis of *F*_*a*_ in the ERP rotates by *ψ*_*r*_ to another fixation axis of *F*. The green ‘primed’ frame in dashed lines shows the green frame after it is rotated at the optical center *O*_*r*_ in the plane containing the vectors ***b***_*r*_ and ***j***_*r*_ that overlays ***b***_*r*_ with ***j***_*r*_. The geometric definition of ocular torsion is the angle 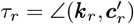.

**Fig 6.**
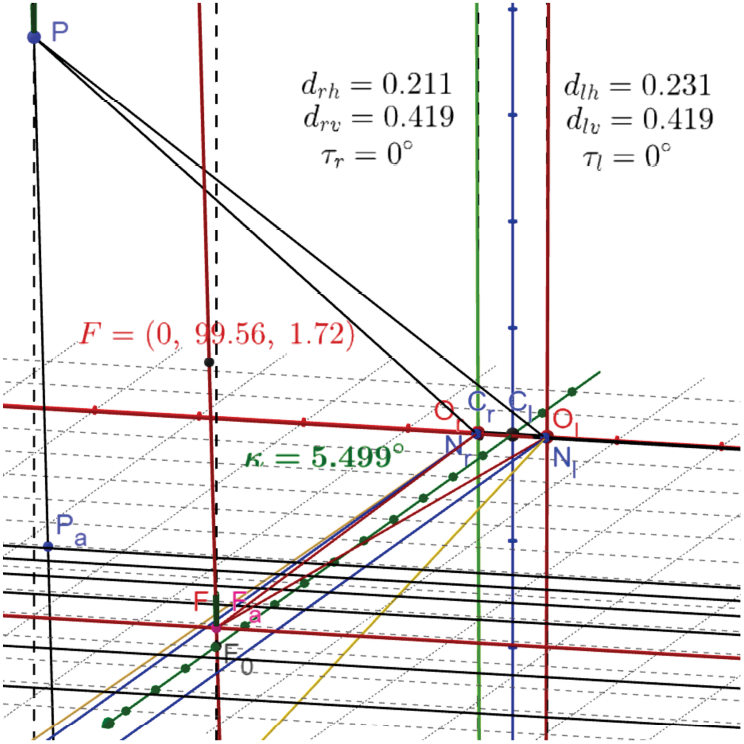
The iso-disparity straight frontal line and vertical horopter in ERP. The fixation *F* = *F*_*a*_(0, 99.56, 1.72) corresponds to the abathic distance fixation of the empirical horopter. The iso-disparity conics are straight frontal lines in the visual plane, and the vertical horopter is tilted by *κ* = 5.5° top away relative to the objective vertical (dashed line). The longitudinal and vertical horopters passing through *F*_*a*_ are shown in red. The point *F*_0_ is the projection of *F*_*a*_ into the head’s transverse plane containing the eyes’ rotation centers (marked with the grid). The other values shown in the figure are explained in the text.

The functional role of the eye’s vertical misalignment of optical components was proposed in [14]. First, by replacing the classic Helmholtz shear theory of ill-defined retinal meridians, which is due to the eye’s misaligned optics, it determines the perceived vertical horopter backward tilt using the criterion of retinal verticality provided by the lens’s vertical tilt. Furthermore, the 3D AE model replaces the Listing plane with the ERP, providing an ab initio binocular version of Listing’s law, which is crucial in oculomotor control. The latter functional role of the eye’s vertical misalignment is examined in detail here.

## Methods

In this section, I discuss the ophthalmological studies on the eye’s resting positions. Then, I review the construction of iso-disparity curves and the definition of ocular torsion in the binocular system with AEs. I conclude this section with an overview of the basic features of *GeoGebra*’s dynamic geometry simulations, utilizing Rodrigues’ vector to visualize transformations when the eyes change binocular positions.

### The eyes’ physiological resting position

Twelve extraocular muscles (six attached to each eye) and a neurological control system work together to control eye position and movement. With the two eyes at rest, both are directed forward in a specific position. The anatomical position of the eyes at rest, observed during electrical silence of the muscles (deep anesthesia or death), is divergent [33]. Estimates of this extent vary from very slight divergence to a divergence angle of about 25 degrees. On the other hand, the physiological position of the eyes at rest maintains a relaxed eye posture, usually with minimally increased energy costs and for prolonged durations without fatigue. This baseline activity level, or tonic (tonus), brings the eyes from the anatomical to the physiological resting position [34].

According to the traditional theories [20, 35], accommodation and vergence adjust to optical infinity at the eyes’ physiological rest posture. Accordingly, oculomotor effort was thought to be necessary only to see near stimuli. Moreover, passive forces (such as the elasticity of supporting tissues) were thought to shift focus and fixation from near to distant stimuli. Helmholtz compared Listing’s Law and the primary position with the minimal energy condition in physics. Contrary to these compelling traditional accounts, the vergence system typically does not adjust to optical infinity (parallel gazes) at rest but assumes an intermediate posture. For example, as early as 1855, Thomas Weber suggested that the eye would focus at an intermediate distance when at ‘complete rest’, see [36].

It has recently become unclear under what conditions we may observe the physiological resting position. The situation is complicated since many factors can influence tonic convergence: fusional convergence, accommodative convergence, proximal convergence, and tonic convergence [37]. Thus, it might be sensible to consider tonic convergence as that convergence in an empty visual field when accommodation is relaxed [38]. In the absence of a stimulus to accommodation, for instance in darkness or a bright but featureless field, the lens is focused not at infinity but at about 1.0 − 1.5 dioptres (lens’ focal length of 1-1.5 meters) [39]. Proximal convergence might also be minimized under these conditions [40].

The tonic vergence of rest measured under degraded stimulus conditions, i.e., with no stimulus for vergence or accommodation, is referred to as the vergence resting position or dark vergence [41, 42]. Dark vergence differs reliably among subjects: the average subject converges at a viewing distance of about 1 m, while the inter-individual range is from infinity to about 40 cm [43, 44]. Moreover, in [43], it was pointed out that distance phoria (eye’s deviation when fusion in binocular viewing is disrupted) is influenced by accommodation for the fixation target. In contrast, dark vergence is a simpler index of tonic vergence [45].

### Iso-disparity curves in binocular system with AEs

Fig. 4 explains for the right eye the construction of iso-disparity conic sections and the vertical horopter using the eyes’ fixation on *F* ^1^(15, 29.56, 7.72) executed from the ERP for the fixation *F*_*a*_(0, 99.56, 1.72).

In this figure, the points *Q*_*r*_ = *O*_*r*_ *± nδ*, where *δ* = 0.005 cm and *n* = 1, 2, …, are shown on the foveal horizon line *H*_*r*_. The *n*-th iso-disparity line is obtained for all pairs (*Q*_*r*_, *Q*_*l*_) such that the difference in distances from *O*_*r*_ and *O*_*l*_ is *nδ*. The subscript ‘*l*’ denotes quantities for the left eye, which are mirror-symmetric in the head midsagittal plane in ERP. For *n* and *n* + 1, the consecutive straight frontal iso-disparity lines for the ERP of fixation point *F*_*a*_(0, 99.56, 1.72) are shown in Fig. 6. Then, the iso-disparity curves are simulated in *GeoGebra* by changing *F*_*a*_ to another fixation point that transforms the iso-disparity lines into ellipses or hyperbolas—Fig. 7 shows the simulation of iso-disparity ellipses for *F*_*a*_ changed to *F* ^1^(15, 29.56, 7.72).

**Fig 7.**
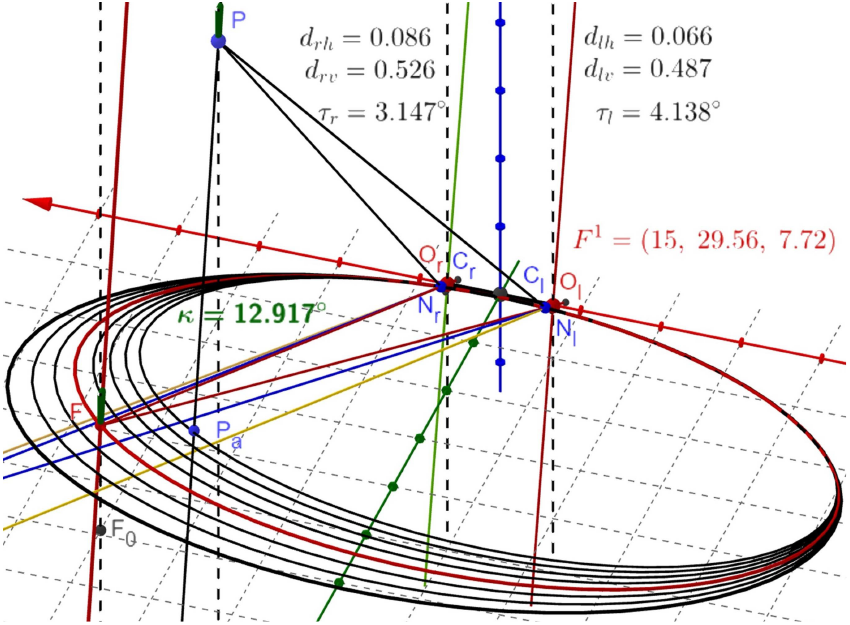
**The iso-disparity ellipses and the vertical horopter for fixation at** *F* ^1^. After the fixation *F*_*a*_(0, 99.56, 1.72) is changed to *F* ^1^(15, 29.56, 7.72), the straight frontal lines change into ellipses, and the vertical horopter changes its tilt to *κ* = 12.92° bottom-away relative to the objective vertical. The values shown here and the calculated disparities are explained in the text.

To construct the subjective vertical horopter, I choose two points *V*_*u*_ and *V*_*d*_ on the apparent vertical line *V*_*r*_ shown in Fig. 4 for the right AE. The points *V*_*u*_ and *V*_*d*_ are the same distance from the optical center *O*_*r*_. The same construction is carried out for the left AE. Two intersections of the projecting rays, one for *V*_*u*_ points and the other for *V*_*d*_ points of the two AEs, define the vertical horopter as explained in [14]. The construction in Fig. 4 resulted in the vertical horopter in Fig. 7 tilted by 12.92° from the true vertical.

### Ocular Torsion and the AE

Ocular torsion is typically described in vision science as a rotation around the visual axis, which represents the direction of the eye’s gaze. For practical purposes, the line of sight connects the point of regard and the center of the entrance pupil, and it is used to indicate the direction of gaze. This axis in the reduced eye model shown in Fig. 1 (a) is identical to the eye’s optical axis. This is not true for the healthy human eye’s misaligned optical components modeled by the AE in Figs. 1 (b) and 3.

The ocular torsion, whose determination is important in clinical diagnosis [27], is often measured using video-oculography as a non-invasive and high-resolution technique. A robust method for ocular torsion measurement using this technique was recently proposed in [46] by tracking the angular shift of the iris pattern around the pupil center, where its advantages over the previous methods were presented.

This work defines the ocular torsion as a rotation around the axis *T*_*r*_ through the optical center *O*_*r*_ parallel to the lens’ optical axis, shown in Fig. 3 for the right eye. From the simulation data in Fig. 3, we obtain |*O*_*r*_*n*_*r*_| = 0.022 cm, indicating that the torsion best approximates the eye’s rolling motion. This ocular torsion definition and its computation are discussed below with the help of Fig. 5.

In the simulation, the black frame (***i***_*r*_, ***j***_*r*_, ***k***_*r*_) is transformed into the green frame (***a***_*r*_, ***b***_*r*_, ***c***_*r*_) when the visual axis throughout *F*_*a*_ in the ERP rotates at *C*_*r*_ to another visual axis throughout *F*. Next, the green frame vector ***b***_*r*_ is rotated at *O*_*r*_ into the black frame vector ***j***_*r*_ in the plane spanned by these two vectors. This rotation is denoted by *ϕ*_*r*_. Because the rotation vector is orthogonal to ***j***_*r*_, it lies in the frontal plane of the ERP (coplanar image planes for the right and left AE). Finally, when this rotation is applied to the green frame, it produces the green dashed frame 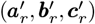 where 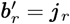. The geometric ocular torsion is 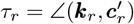.

However, when the AE is rotated about *C*_*r*_, the frame vectors are rotated and translated as they move with the image plane. The rotation angle between vectors is unaffected by translation. For simplicity, in Fig. 5, all frame vectors are shown at *O*_*r*_ in the ERP. This is justified by the fact that the translation is about that 0.02 cm for typical eye rotations, using the simulation data in Fig. 3; cf. [14]. Later, in simulations of ocular torsion, when the visual axes pass through their respective optical centers, they are always translated accordingly with the change in the fixation point.

Because the AEs in the ERP are mirror-symmetric relative to the head’s midsagittal plane, the above discussion establishes the geometrical calculations of the eyes’ ocular torsion for any binocular fixation executed from the ERP. The actual simulation of ocular torsion of each eye in the binocular system with EAs is given in Figs. 8 and 9.

**Fig 8.**
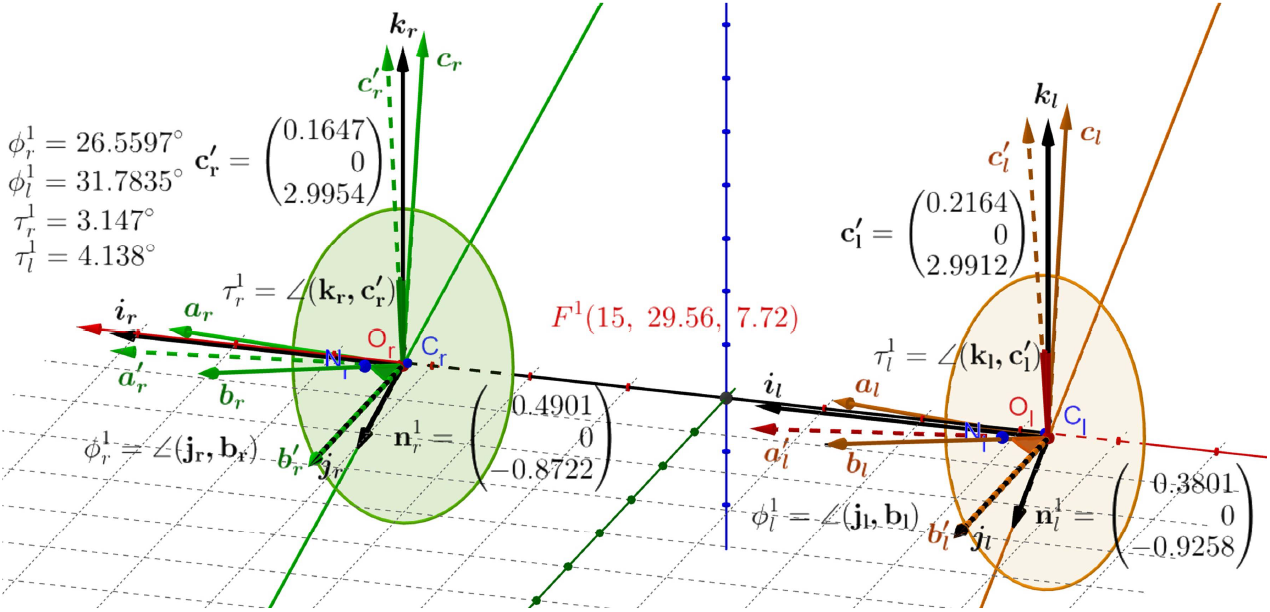
**Ocular torsion when** *F*_*a*_ *→ F* ^1^. The black frames in the ERP (***i***_*r*_, ***j***_*r*_, ***k***_*r*_) and (***i***_*l*_, ***j***_*l*_, ***k***_*l*_) for the right and left AEs are attached at the image planes’ optical centers *O*_*r*_ and *O*_*l*_. Their orientations agree with the upright head’s coordinate system. Thesolid-colored frames are moving from the initial position of the black frames when fixation *F*_*a*_ changes to fixation *F* ^1^. The dashed-colored frames are obtained by rotating the solid-colored frames by moving ***b***s vectors onto ***j***s vectors. The vectors 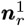 and 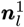 give the rotation axes. They are contained in the co-planar image planes of the ERP, parallel to the head frontal plane.

**Fig 9.**
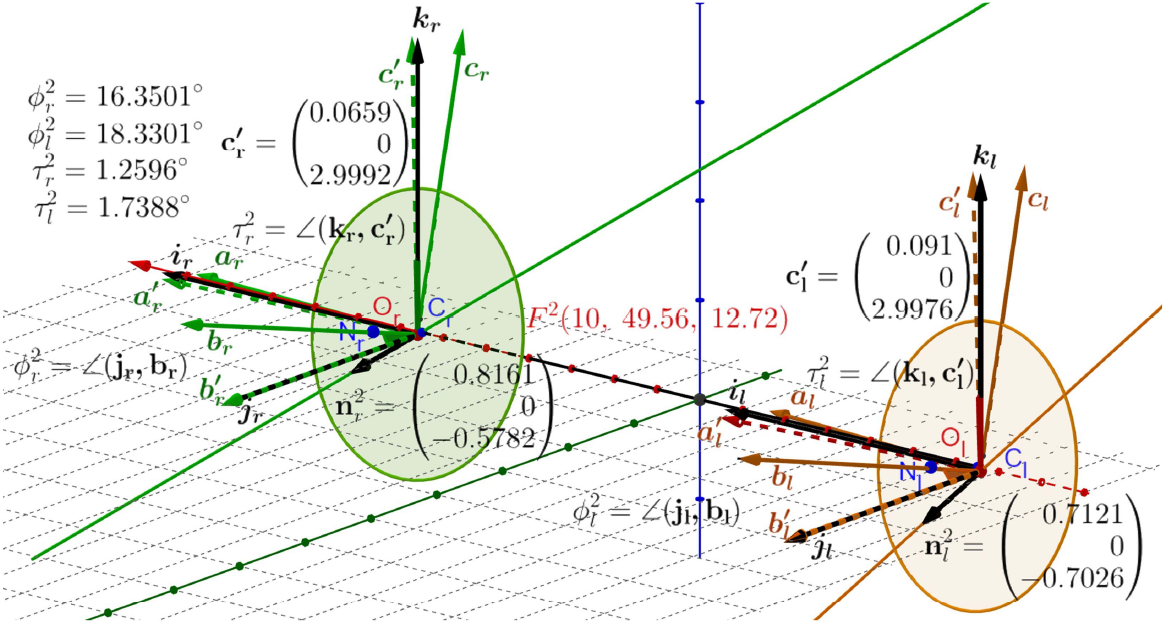
**Ocular torsion simulation when** *F*_*a*_ → *F* ^2^. For the fixation *F* ^2^(10, 49.56, 12.72, the frames’ angles of rotations around the axes 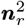 and 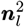 and ocular torsion of each eye are given in the figure.

I note that for a single symmetric eye (all angles of asymmetry in AE are zero and *O*_*r*_ agrees with *C*_*r*_), the rotation from the primary position into secondary positions is torsion-free, i.e.,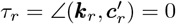. However, when the human eye’s asymmetry of optical components is taken into account, ocular torsion is zero at the ERP. Moreover, it remains zero after vertical movement from the ERP of binocularly fixating eyes, but is non-zero after a horizontal movement, see [14] for discussion.

### *GeoGebra* Simulations and Rodrigues’ vector

The iso-disparity conic sections, the vertical horopter, and ocular torsion are computed and visualized in *GeoGebra*’s dynamic geometry simulations in the binocular system in which the AE models the human eye’s 3D misalignments of the fovea and the lens. The simulated iso-disparity conic sections in the visual plane and the vertical horopter in Figs. 6 and 7, as well as, ocular torsion in Figs. 8 and 9 and the half angle rule in Figs. 10 and 11, are generated by an input into *GeoGebra*, changing the initial ERP’s fixation *F*_*a*_(0, 99.56, 1.72) coordinates. For simulations with *GeoGebra* for different fixations, see [12–14].

**Fig 10.**
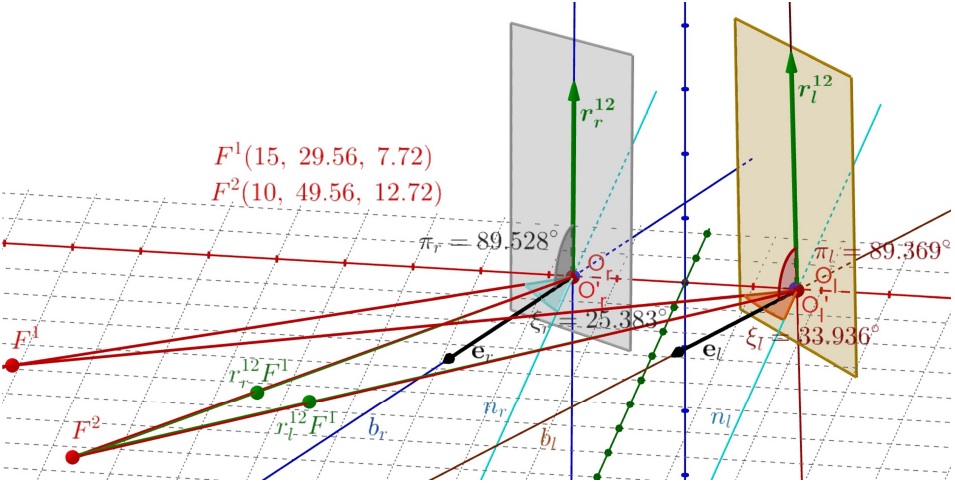
**The half-angle rule for** *F* ^1^ *→ F* ^2^. The green dots 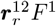 and 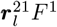 are the rotations of *F* ^1^ by the indicated Rodrigues’ vectors. The green lines through the green dots and 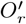 and 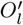 overlying the visual axes of *F* ^2^ are obtained by these Rodrigues’ vectors; they generate the eyes’ rotations from *F* ^1^ to *F* ^2^. The vectors ***e***_*r*_ and ***e***_*l*_ are bisectors between lines *n*_*r*_ and *n*_*l*_ perpendicular to the ERP and the visual axes of *F* ^1^ for the left and right AE. The displacement planes are perpendicular to the bisectors.

**Fig 11.**
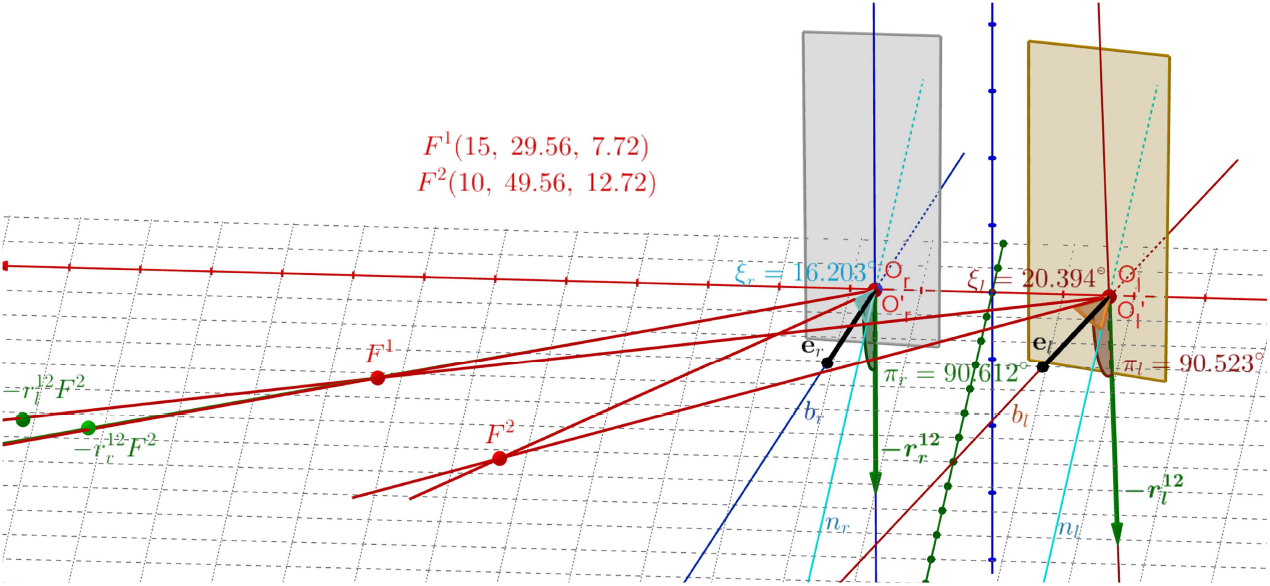
**The half-angle rule for** *F* ^2^ *→ F* ^1^. The green dots 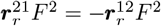 and 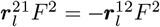 are the rotations of *F* ^2^ by the indicated Rodrigues’ vectors. The green lines through the green dots and *O*_*r*_ and *O*_*l*_ are overlying the visual axes of *F* ^1^ for the right and left eye. Notation is similar as in Fig. 10.

*GeoGebra* simulations of the eyes’ binocular postures and related retinal coordinates in space (iso-disparity conic sections), include geometric primitives (lines, conics, planes, spheres, cylinders, and the like), geometric constructs (parallel lines, perpendicular lines to planes, bisectors, and the like), and basic geometric transformations (translations, rotations, reflections, and the like). These geometric tools provided in *GeoGebra* are used directly to design the visualization, so programming is not involved.

However, the primary challenge in creating a dynamic simulation for this study is connecting the geometric tools. It simulates the iso-disparity conics, the vertical horopter, and ocular torsion when the ocular system with AEs changes under the binocularly constrained eyes’ rotations by moving the fixation point, visualizing their 3D transformations. In the simulations presented in this study, the number of geometric tools increases rapidly, reaching over 100 objects, while those that create connections are hidden. It prevents an easy understanding of the simulation construction. Moreover, a small unintended change in any of those tools and their connections can disable the simulation.

Because the *GeoGebra* does not involve programming, the links to simulations of ocular torsion in the binocular system with AEs are available with the author’s support upon reasonable requests.

In *GeoGebra* simulations of eyes’ positions, all rotations in 3D are performed with Rodrigues’ vector as it encodes each rotation axis and angle. It also allows the composition of eye rotations, which is particularly important in the binocular half-angle rule simulation. To introduce Rodrigues’ vector framework, I start with the conclusion from Euler’s rotation theorem: Any rotation matrix *R* can be parametrized as *R*(*ϕ*, ***n***) for a rotation angle *ϕ* around the axis with the unit vector ***n***. This parametrization is unique if the orientation of 0 < *ϕ* < 180° is fixed. Usually, a counterclockwise (or right-hand) orientation is chosen to get angles’ positive values.

The pair *ρ* = cos(*ϕ/*2), ***e*** = sin(*ϕ/*2)***n*** is known as *Euler-Rodrigues parameters* and

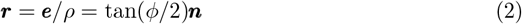

as *Rodrigues’ vector*, usually referred to as *rotation vector* [47]. Rodrigues proved that under composition *R*^*′′*^(*ϕ*^*′′*^, ***n***^*′′*^) = *R*^*′*^(*ϕ*^*′*^, ***n***^*′*^)*R*(*ϕ*, ***n***), the corresponding Euler-Rodrigues parameters transform as follows [48],

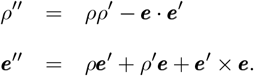

Then, setting ***r***^*′′*^ = ***e***^*′′*^*/ρ*^*′′*^, we see that the composition for Rodrigues’ vectors is given by

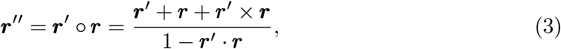

Further, ***r***^−1^ = −***r*** and tan(*−ϕ/*2)***n*** = tan(*ϕ/*2)(−***n***) define the same rotation vector.

Also, one Rodrigues’ vector corresponds to one rotation, and the Rodrigues’ vector corresponding to the unit matrix is the zero vector [47].

Note that (cos(*ϕ/*2), sin(*ϕ/*2)***n***) is a unit quaternion that describes the rotation by *ϕ* around ***n***. Quaternions were introduced in 1957 by Westheimer [49] to describe eye kinematics. Quaternions and Rodrigues’ (rotation) vectors have often been used to analyze the eye’s rotations with the connection to Listing’s law, cf. [50–53].

## Results

### Listing’s law and half-angle rule modifications for AEs

The eyes’ tonic vergence of rest is measured under degraded stimulus conditions, i.e., with no stimulus for vergence and accommodation. This measurement differs reliably among subjects: a typical subject converges at a viewing distance of about 1 m, while the inter-individual range is from infinity to about 40 cm [43, 44]. The estimation of the range of ERP’s fixation point *F*_*a*_ in [11] was as follows: using the average values of the interocular distance 2*a* = 6.5 cm, the angles *α* = 5.2° and −0.4° ≤ *β* ≤ 4.7° in Eg. 1, we obtain 34 ≤ |*OF*_*a*_| ≤ 380 in centimeters with the average of about 1 meter for the typical value *β* = 3.3°. It was also noted there that the distance in Eq. 1 increases to infinity (when the value of *β* approaches the value of *α*), as occasionally reported in experimental measurements.

In a 3D setting of the binocular system with AEs, the image planes for the ERP binocular posture are coplanar and constitute the frontal plane of the stationary upright head. Given the above discussion of changes in the tonic vergence rest corresponding to changes in ERP for typical values of misalignment angles in the human population, I substitute the ERP’s frontal plane for the Listing plane. The cyclopean gaze direction with the asymmetry angles *α* = 5.2°, *β* = 3.3°, *γ* = −2°, and *ε* = −1° in this posture passes through the point *F*_*a*_(0, 99.56, 1.72), expressed in centimeters, and is not perpendicular to the frontal plane, so strictly speaking this ERP’s plane is not the Listing plane. It has ophthalmological underpinnings, and, according to standard terminology, it can be referred to as the primary displacement plane. The angle value between the substitutions for the Listing plane and the primary position differs from 90° by about 1°. Of course, the gaze of each eye differs more from perpendicularity; it can be estimated from data in Fig. 3 to be about 1.8°.

Further, following the calculations presented in Table 2 in [14], for the AE angular parameters *α* = 5.2°, *β* = 3.3°, *γ* = −1°, and *ε* = −1° and for parameters *α* = 5.2°, *β* = 3.3°, *γ* = −2°, and *ε* = −2° the ERP fixation is *F*_*a*_(0, 99.61, 0) and the deviation from perpendicularity vanishes whereas for angular parameters *α* = 5.2°, *β* = 3.3°, *γ* = −3°, and *ε* = −1° the ERP fixation is *F*_*a*_(0, 99.55, 3.45) and the deviation from perpendicularity is about 2°.

The above discussion of the angle between the ‘cyclopean gaze’ direction through the fixation point *F*_*a*_ to the ERP frontal plane for different values of optical component misalignments in AE can be compared, although not directly, to Listing’s law validity measurements only for a single eye. The thickness of the Listing plane, which indicates the precision of Listing’s law, as reported in the literature, is as follows: [15] found a mean thickness of 1.5° for fixations, whereas [54] reported 1.4 *±* 0.5°. I note that from the data in Fig. 2, the angle between the visual axis of the right eye and the line perpendicular to the ERP is about 2.15° for the eye’s misalignment considered in this study.

The binocular ERP not only provides the necessary neurophysiological meaning for the primary position and Listing plane but also supersedes the original monocular Listing’s law with the ab initio formulated binocular law. This is significant because, as pointed out in [21], the oculomotor system couples the 3D rotations of the eyes and, therefore, their torsion. On the other hand, Listing’s law constrains the eye’s redundant torsional degree of freedom during the eye’s changing position to facilitate spatial understanding [20].

Tweed and Vilis in [51] pointed out that when Listing’s law and the half-angle rule are applied to studies of smooth pursuit and saccadic eye movement, because the Rodrigues’ vector is not the time-integral of angular velocity, the mathematical framework of 3D rotational kinematics should be employed. However, when considering the transformation between two stationary postures, the angular velocity vector is not well defined. Later, the binocular half-angle rule is tested to high accuracy between two tertiary positions in Rodrigues’ vector framework, and even despite the eye’s misaligned optical components included.

### Simulations of iso-disparity conic sections and vertical horopter

Two simulations of the iso-disparity conic sections and the vertical horopter, one in the ERP of the abathic fixation *F*_*a*_(0, 99.56, 1.72) and the other for the tertiary posture after the fixation point *F*_*a*_ changed to *F* ^1^(15, 29.56, 7.72). They are shown in Fig. 6 and Fig. 7, respectively. These two examples of simulations are used later in ocular torsion calculations and discussions of the binocular half-angle rule, which is the extension of Listing’s law to eye rotations from tertiary postures.

The iso-disparity conic sections in Fig. 6 are straight frontal lines in the ERP, which, as discussed before, hold for the fixation *F*_*a*_ distinguished by coplanar image planes parallel to coplanar lens’ equatorial planes. Additionally, in this posture, the vertical horopter is a straight line tilted from the true vertical by *κ* = 5.5°, with its top positioned away from the head. This value is observed in experiments, [55, 56]. From the data of projections of point *P* in the binocular field into the image planes of the corresponding right and left AEs, the horizontal disparity for the right and left AEs is *δ*(*P*) = *d*_*lh*_ *− d*_*rh*_ = 0.02 cm. The vertical disparity and the torsional disparity are zero.

Fig. 7 shows the iso-disparity ellipses and the vertical horopter for the tertiary fixation *F* ^1^(15, 29.56, 7.72). The horizontal disparity value is *δ*(*P*)_*h*_ = 0.02 cm, the vertical disparity value is *δ*(*P*)_*v*_ = 0.039 cm and the torsional disparity value is *δ*_*t*_ = *τ*_*r*_ *− τ*_*l*_ = −0.99°. The vertical horopter is tilted 12.92° bottom-away from the true vertical, which has never been tested in experiments.

In both figures, the projection of *P* into the visual plane along the parallel line to the vertical horopter, *P*_*a*_, is exactly on the fourth consecutive disparity line down from the horopter (red frontal line through *F*_*a*_). Because the horizontal retinal relative disparity value between each consecutive disparity line is 0.005 cm for the simulations (cf. Fig. 4), the spatial disparity of *P* (*P* is different in each figure) agrees with the retinal horizontal disparity in both Figs. 6 and 7,

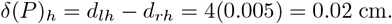

This result indicates that the subjective vertical horopter agrees with the perceived vertical direction, as reported in [56].

The computation and visualization of ocular torsion shown in Fig. 7 are presented in the next section. As mentioned above, due to the misalignment of the optical components of the eyes, the ocular torsion for rotations from ERP is generally nonzero. It is zero at the ERP, and when the eyes change the fixation from *F*_*a*_ to the vertically displaced *F*, the ocular torsion remains zero. I refer the reader to [14] for a detailed discussion of results for basic 3D rotations from *F*_*a*_.

### Binocular torsion for rotations from the ERP

Before reading this section, the reader should recall the definition of ocular torsion explained in Fig. 5. The binocular torsion for each eye is computed for rotations from the ERP; the binocular substitution for the original Listing’s plane of a single eye. It is simulated in *GeoGebra* and shown in Fig. 8 for the change of fixation *F*_*a*_(0, 99.56, 1.72) of the ERP to tertiary posture with the fixation at *F* ^1^(15, 29.56, 7.72). The data from the simulation presented in this figure supports all assertions made in the discussion.

In Fig. 8, when *F*_*a*_ changes to *F* ^1^, the solid green frame for the right AE, (***a***_*r*_, ***b***_*r*_, ***c***_*r*_), and the brown frame for the left AE, (***a***_*l*_, ***b***_*l*_, ***c***_*l*_), are moving with the image planes from their position that initially in ERP agrees with the black frames, (***i***_*r*_, ***j***_*r*_, ***k***_*r*_) and (***i***_*l*_, ***j***_*l*_, ***k***_*l*_), for the respective AE. From now on, when writing Rodrigues vectors, the associated rotation angles and unit vectors, I use superscript notation, such as 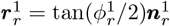, which is Rodrigues’ vector for the right eye.

To calculate each eye’s ocular torsion, the green and brown frames are rotated such that ***b***s vectors are overlaid with ***j***s vectors. These rotations are about the optical centers in the planes spanned by ***b***s and ***j***s vectors for each eye such that the unit vectors 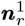 and 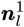of the rotation axes are perpendicular to these planes and, therefore, contained in the coplanar image planes of the ERP. The rotations angles are 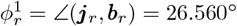 and 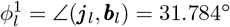. When applied to these colored frames, these rotations result in two ‘primed’ frames in dashed lines, green frame 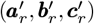, and the brown frame 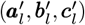. These last rotations lead to the definition of ocular torsion and its value for each eye, 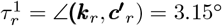 and 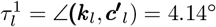. Thus, the torsional disparity 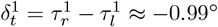. All counterclockwise right-hand rotations are positive.

In the *GeoGebra* simulation in Fig. 8, Rodrigues’ vectors (rotating the frame vectors ***j*** s onto ***b***s) in the binocular system with AEs,

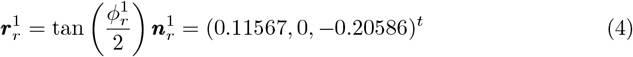

and

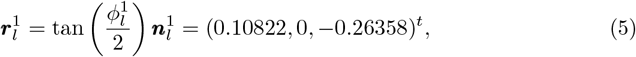

for the right and left eye, respectively. The superscript “t” is the transpose, making column vectors. These vectors efficiently rotate geometric objects in *GeoGebra*’s simulations,Similarly as before, from the data in Fig. 9 when *F*_*a*_(0, 99.56, 1.72) changes to *F* ^2^(10, 49.56, 12.72), Rodrigues’ vectors are

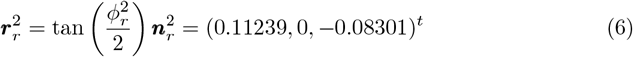

and

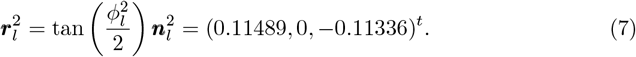

The data from the simulation in Fig. 9 again allows the calculation of each eye’s ocular torsion and the torsional disparity for the fixation *F* ^2^(10, 49.56, 12.72).

### Binocular half-angle rule

The binocular half-angle rule is next constructed and simulated between the tertiary positions when the eyes change from *F* ^1^(15, 29.56, 7.72) to *F* ^2^(10, 49.56, 12.72). From the definitions of Rodrigues’ vectors in Eq. 2 and their composition Eq. 3, the sequence of the changes in the fixations *F*_*a*_ *→ F* ^1^ *→ F* ^2^ results in the composition of Rodrigues vectors for the respective eyes:

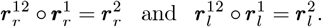

Here, the Rodrigues’ vectors 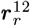 and 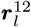 correspond to the right and left eye’s rotations when bifoveal fixation *F* ^1^ changes to *F* ^2^ in the stationary upright head. Also, Rodrigues’ vectors 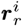 and 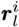 correspond to the change *F*_*a*_ *→ F* ^*i*^ where *F*_*a*_ is fixation of the ERP and *i* = 1, 2.

Using that ***r***^−1^ = −***r***, the above equations written as follows:

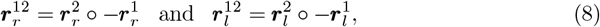

are compositions of Rodrigues’ vectors for the right and the left eye corresponding to the composition of rotations 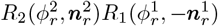 and 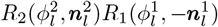 when fixation changes from *F* ^1^(15, 29.56, 7.72) and *F* ^2^(10, 49.56, 12.72).

Using Eqs. 3–8 and calculating the cross and scalar products, Rodrigues’ vectors 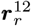 and 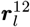 are the following:

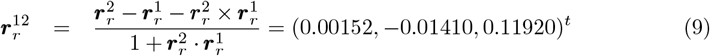

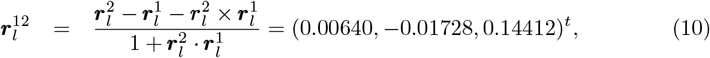

Next, using Eqs. 2, 9, and 10, we have

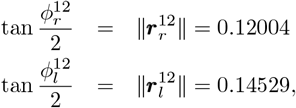

which finally gives the corresponding angle values 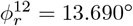 and 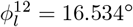 for the right and left eye, respectively.

We immediately recognize that the sequence of the changes in the fixations *F*_*a*_ *→ F* ^2^ *→ F* ^1^ results in the composition of Rodrigues vectors for the respective eyes:

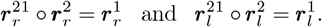

and

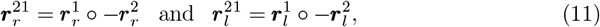

are compositions of Rodrigues’ vectors for the right and the left eye corresponding to the composition of rotations 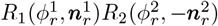 and 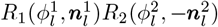 when fixation changes from *F* ^2^(10, 49.56, 12.72) to *F* ^1^(15, 29.56, 7.72).

Further, following similar lines as before to obtain Eqs. 9 and 10,

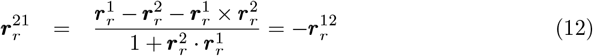

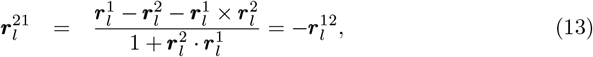

To my knowledge, it is the first time that the half-angle rule obeying a simple noncommutative rule has been demonstrated to uphold the binocularly constrained fixations in a stationary upright head. Moreover, it is a general rule that also applies to the binocular system with misaligned eyes’ optics. Usually, the half-angle rule has been related to the ‘effect’ of noncommutativity of rotations, which has never been precisely formulated. The above discussion of the binocular half-angle rule is next demonstrated in *GeoGebra* simulations.

In the first simulation in Fig. 10, the fixation changes between tertiary positions from *F* ^1^ to *F* ^2^. The displacement planes containing Rodrigues’ vectors are determined by the bisectors between the visual axes of *F* ^1^ and the perpendicular lines *n*_*r*_ and *n*_*l*_ to the ERP plane at the optical centers *O*_*r*_ and *O*_*l*_ for the right and left eye, respectively, cf. Fig. 10. This figure shows the simulation, which differs from the one used in [14].

In the second simulation in Fig. 11, the fixation changes between tertiary positions from *F* ^2^ to *F* ^1^. The displacement planes containing Rodrigues’ vectors are determined by the bisectors between the visual axes of *F* ^2^ and the perpendicular lines *n*_*r*_ and *n*_*l*_ to the ERP plane at the optical centers 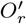 and 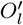 for the ight and left eye, respectively, cf. Fig. 11.

The simulation tests whether the computed Rodrigues’ vectors 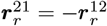 and 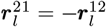 lie in the corresponding displacement planes determined by the bisectors. It also verifies that these Rodrigues’ vectors generate the fixation *F* ^2^ by using them to rotate *F* ^1^; the lines through the green dots 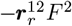 and 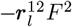 and the points 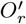 and 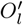 perfectly overlying the red visual axes.

The displacement planes (shown as rectangular areas) are defined for the respective eyes by the vectors ***e***_*r*_ and ***e***_*l*_ along the bisectors. The angles between 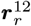 and ***e***_*r*_ has the value of *π*_*r*_ = 89.53° and between 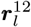 and ***e***_*l*_ the value of *π*_*l*_ = 89.37° in the first simulation and the angles between 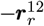 and ***e***_*r*_ has the value of *π*_*r*_ = 90.61° and between 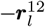 and ***e***_*l*_ the value of *π*_*l*_ = 90.52° in the second simulation. It shows the approximations of the half-angle rule for each eye, due to the misalignment of the eye’s optical components.

### The binocular configuration space

The eyes’ configuration specifies their stationary posture of bifoveal fixation in the binocular system of a stationary upright head. The element of the configuration space (CS) for each eye comprises pairs of torsion-free rotation of the change in the visual axis direction described by Rodrigues’ vectors and the change of ocular torsion after this rotation. I recall that Rodrigues’ vector has three independent parameters, the minimal number needed to specify the rotation, and one Rodrigues’ vector corresponds to one rotation matrix. Thus, it is particularly well-suited to formulate the CS.

Most significantly, this CS, supported by the ophthalmology studies and simulations, naturally extends Listing’s law with the eyes rotating from or to the ERP and rotations between tertiary positions obeying the half-angle rule and satisfying a simple noncommutative rule compatible with the constraints of eyes’ bifoveal fixations.

Before I formulate the CS, I need to discuss how ocular torsion of each eye, explained in Fig. 5 and simulated in Figs. 8 and 9 in *GeoGebra*, can be computed. Without specifying the eye in the binocular system, Rodrigues’ vector when the fixation axis given by vector 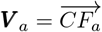 changes to the visual axis given by vector 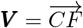 can be computed as follows. The angle between the fixation axes is

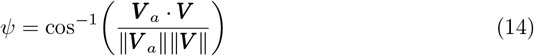

such that Rodrigues’ vector is ***Q*** = tan(*ψ/*2)***N*** where

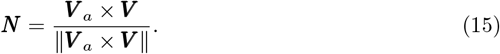

is the unit vector of the rotation axis. Rodrigues vectors ***Q***_*r*_ for the right eye and ***Q***_*l*_ for the left eye define the corresponding rotation matrices *R*(*ψ*_*r*_, ***N*** _*r*_) and *R*(*ψ*_*l*_, ***N*** _*l*_) [47].

In Fig. 8 for fixation point *F* ^1^, the Rodrigues’ vectors ***Q***_*r*_ and ***Q***_*l*_ are used to rotate the corresponding initial frame vectors (***i***_*r*_, ***j***_*r*_, ***k***_*r*_) and (***i***_*l*_, ***j***_*l*_, ***k***_*l*_) into the frame vectors (***a***_*r*_, ***b***_*r*_, ***c***_*r*_) and (***a***_*l*_, ***b***_*l*_, ***c***_*l*_), denoted in green for the right eye and in brown for the left eye.

Further, rotation by Rodrigues’ vectors 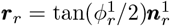 and 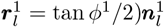, first defined by rotating ***b***_*r*_ back onto ***j***_*r*_ and ***b***_*l*_ back onto ***j***_*l*_, are applied with opposite rotation angles to the frame vector ***c***_*r*_ and ***c***_*l*_, giving the frame vectors 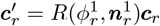 and 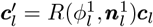, respectively. Then, ocular torsion is 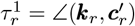 of the right eye and 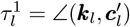 for the left eye.

I unify the notation to account for all eye rotations between *F*_*a*_ and *F* ^*k*^ and rotations between *F* ^*k*^ and *F* ^*l*^. To do this, I change *F*_*a*_ to *F* ^0^ and denote ***r***^1^ by ***r***^01^ when *F* ^0^ *→ F* ^1^. It allows to write ***r***^*kl*^ when *F* ^*k*^ *→ F* ^*l*^, *k* = 0, 1, 2, *l* = 0, 1, 2. Then, ***r***^*lk*^ = −***r***^*kl*^ when *F* ^*l*^ *→ F* ^*k*^. For changes in ocular torsion in these cases, I use the notation *τ* ^*kl*^ *≡ τ* ^*k*^ *→ τ* ^*l*^. I also write *τ* ^*lk*^ = *−τ* ^*kl*^. I recall that *τ* ^0^ = 0°. With this notation, the elements of the CS are pairs 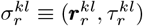 for the right eye and 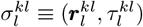 for the left eye.

Without specifying the eye in the binocular system with AEs, the binocularly constrained eyes’ changes involving fixations *F* ^0^, *F* ^*k*^, *F* ^*l*^, and *F* ^*m*^ are the following:

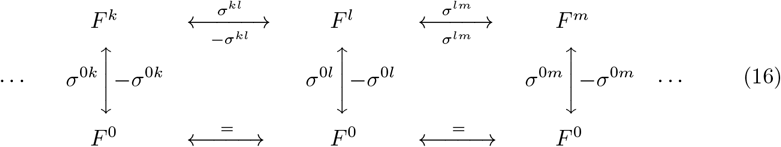

where the minus sign indicates a down arrow or a left arrow.

For the composition of the CS elements, I need only discuss the composition of Rodrigues’ vectors. It is associative, for example,

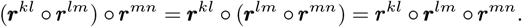

It is noncommutative, but with a simple noncommutativity rule, for example,

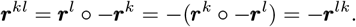

Note that ***r***^*kl*^ transforms the fixation from *F* ^*k*^ to *F* ^*l*^ and −***r***^*kl*^ transforms from *F* ^*l*^ to *F* ^*k*^, cf. Figs. 10 and 11.

Lets examine the path: *F* ^*m*^ *→ F* ^0^ *→ F* ^*l*^ *→ F* ^*k*^. The composition of the corresponding Rodrigues’ vectors is

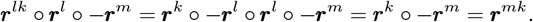

Recall that the composition order of Rodrigues’ vectors is the same as the composition order of the corresponding rotation matrices, i.e., from right to left.

## Discussion

The results in this study are obtained based on geometric considerations without any input from experiments or neural circuits modeling, which distinguishes it from typical publications in the field of vision science. However, this study not only employs an abstract geometric eye model but also incorporates the anatomy and physiology of the eye, thereby supporting the biologically mediated aspects of binocular vision that it aims to model. This aspect also distinguishes it from psychophysical lines of inquiry [57].

In the geometric model of the binocular system with AEs, ocular torsion constrained by bifoveal fixations is computed in Rodrigues’ vector framework and visualized in *GeoGebra* dynamic geometry simulations. It requires that in Listing’s law, originally formulated for a single eye, the Listing plane is replaced with the ERP corresponding to the empirical horopter’s abathic distance fixation. The ERP is binocular and corresponds to the eyes’ resting vergence, in which the eye muscles’ natural tonus resting position serves as a zero-reference level for convergence effort. Supported by ophthalmology studies, the replacement of the Listing plane with the ERP provides the physiologically underpinned ab initio binocular reformulation of Listing’s law.

This replacement of the Listing plane with the ERP revises the elusive neurological significance of the primary position of Listing’s law [58]. It is corroborated by many attempts to define the primary eye position [20, 59, 60]. Moreover, none of those definitions can be precisely satisfied with the human eye’s misaligned optical components. For example, for horizontal changes in binocular fixation executed from the ERP, as opposed to vertical changes, the ocular torsion has a nonzero value; however, it is accompanied by a small torsional disparity. More importantly, the straight frontal iso-disparity lines in the ERP change into hyperbolas under the horizontal rotation of the eyes. It can affect our impression of being immersed in the ambient environment, a remarkable feat considering our eyes receive 2D projections. The reader is directed to [13, 14] for a comprehensive discussion.

The need for an unambiguous binocular formulation of Listing’s law was mentioned, for example, in [21]. The ad hoc binocular modification of Listing’s law by tilting the Listing planes by angles proportional to vergence-dependent coefficients has been a subject of controversy. Moreover, a binocular control system couples the 3D movements of the eyes and, hence, couples ocular torsion of the fixating eyes. The repercussions included numerous experimental inconsistencies [21–23]. Consequently, as pointed out in [21], there is no generally accepted explanation for Listing’s law, which remains true today.

The half-angle rule states that for Listing’s law to be obeyed, the Listing plane must tilt by half the angle of eye eccentricity from the primary position. It was later attributed to the noncommutativity of 3D rotations [61]. The most often expressed purpose of Listing’s law and the related half-angle rule is to assist the brain in controlling the eyes’ torsional positions, thereby restricting the eyes’ movement from 3D to 2D and, hence, minimizing the noncommutativity problem. Although general rotations are noncommutative, how this noncommutativity relates to bifoveal fixations was never clarified before the results presented here in Figs 10 and 11.

An alternative point of view, advocated in [62, 63], posits that the brain makes no special effort to constrain the torsional position of the eye to Listing’s plane. It is supported by a fortunate arrangement of extraocular muscles and orbital connective tissues, which makes it appear commutative to the brain [64]. See also review in [65]. In the studies presented here, the computation of ocular torsion in binocular fixations is a consequence of Euler’s rotation theorem implemented in the framework of Rodrigues’ vector. There is no special effort to satisfy any preset conditions to constrain the torsion position other than binocular constraints. In this way, this work aligns with the aforementioned alternative perspective.

Further, in the geometric study presented here, the binocular half-angle rule was constructed and tested in simulation for changes between two tertiary positions using the composition of Rodrigues’ vectors. It was surprisingly very well satisfied for each eye during binocular fixations, even with the included eye’s misaligned optical components. Of course, by looking around in near space, the brain’s priority is to maintain bifoveal fixations as precisely as possible. Nevertheless, it is surprising that the half-angle rule supports the constraints of the 3D binocular eyes’ movements with a fixed upright head. It is reasonable to expect that during the eyes and head movements, the geometric bifoveal constraints should include noncommutativity control by the brain’s neural circuits.

The simulation of the half-angle rule, which were obtained here for tertiary postures when the fixation changed from *F* ^1^(15, 29.56, 7.72) to *F* ^2^(10, 49.56, 12.72), allow us to consider two different descriptions of a sequence of eyes’ binocular posture execution in a stationary upright head. I again need only to discuss the Rodrigues’ vectors. In the first description, the change between the two consecutive postures is directly related to the preceding and the following postures, as shown in the following expression:

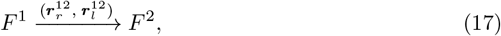

where 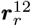 and 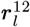 are Rodrigues’ vectors (9) and (10). The first execution in Rodrigues’ vector framework yielded the binocular half-angle rule between two tertiary postures, as shown in the simulation in Figure 10.

On the other hand, in the second description, changes between any two consecutive postures are related to the ERP for fixation *F*_*a*_,

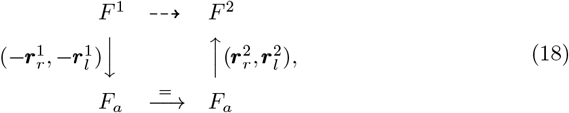

where 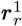 and 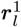 are Rodrigues’ vectors in Eqs. 4 and 5 for fixation *F* ^1^ and 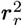 and 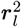 in Eqs. 6 and 7 for fixation *F* ^2^. This is a commutative diagram in which the dashed arrow indicates the composition of transformations given by continuous arrows. From the discussion of the previous section, the expressions in Eqs. 17 and 18 can be reversed by changing the signs of Rodrigues’ vectors and directions of arrows.

The second description of the consecutive eye binocular posture shifts relative to the ERP may be more straightforward and could reveal how the visual system functions. It seems to be supported by experimental observations. In [66], the experimental measurements supported by photographic and video analysis demonstrated that the anatomically determined primary position is a natural constant position to which the eyes are automatically reset from any displacement of the visual line. Further, the evidence indicated an active neurologic basis for the primary position.

The results of the construction and simulations of the half-angle rule in Figs. 10 and 11, along with a detailed discussion of the composition of rotations in terms of Rodrigues’ vectors, provide a framework for constructing the configuration space of binocularly constrained discrete fixations. It is important because a large part of our lives is in confined spaces, and we must perceive with bifoveal fixations on near objects. For continuous eye rotations, however, specifying the configuration space of the binocularly fixating eyes as a subset of *SO*(3) *× SO*(3) or the formulation in terms of the angular velocity vectors is a challenging problem. The configuration space, on the other hand, is simple for Hering’s ‘double’ eye, characterized by conjugate eye movements with both eyes having parallel lines of sight. In such cases, the configuration space is the set of curves in *SO*(3), as seen in [67, 68], for example.

The half-angle can only be satisfied approximately for the misaligned eyes’ optical components; its simulation results should be compared with the experimental data. However, most experiments testing Listing’s law and the half-angle rule were performed for pursuits and saccades, and only a few for fixations. The systematic deviations from Listing’s law were demonstrated in [69], which involved four healthy subjects using the coil mounted on the viewing eye. The appropriately designed study should investigate how these systematic deviations depend on the eye’s angles of optical misalignment.

Although only fixations are considered here, the experimental findings in [70–73] show that Listing’s law and the half-angle rule are not perfectly satisfied during human saccades or pursuit movements. Nevertheless, the measurements of human eye movements in [74] show, on the other hand, that Listing’s law holds with remarkable accuracy during smooth pursuit.

Listing’s law and the half-angle rule are biologically implemented by kinematic constraints that limit the range of eye positions and angular velocities used by the eyes. These constraints have been attributed to relative contributions of the neurally generated commands and the physical mechanics of the eye and its surrounding muscles and tissues [64, 65, 75–79]. These aspects were recently tested experimentally in [80] and [81], in both studies for Rhesus monkeys. The results indicated that, despite a clear role for the brain in coordinating movement kinematics, the ocular plant makes an important peripheral contribution to mechanically optimizing the kinematic constraints necessary for visually guided eye movements. However, the half-angle rule cannot be satisfied accurately due to the human eye’s misaligned optical components. The potential impact of the eye’s misaligned optics on the oculomotor system coordinating the eyes’ movements should be investigated.

Finally, I will comment on the literature discussing the relevance of Listing’s law to clinical practices. In [82], it is stated that acute fourth nerve palsy violates Listing’s law during saccades. In contrast, chronic palsy obeys it, indicating that neural adaptation can restore Listing’s law by adjusting the innervations to the remaining extraocular muscles, even when one eye’s muscle remains partially paralyzed. The use of Listing’s law was clinically evaluated in patients with strabismus [22, 83]. Although the study in [22] involved one patient and no general conclusion was reached, the study in [83] made specific recommendations. In [83], the author first notes that the clinical evaluation of strabismus has traditionally focused on horizontal and vertical alignment, without providing information about torsion and stereo vision. In those studies, some questions remained unanswered: What are the effects of strabismus surgery on 3D eye orientation, or what are the consequences of surgery for torsion despite good eye realignment? Since the answers involve binocular Listing’s law, which was only precisely formulated in the study presented here, the results of this study should be a part of those answers.

## Availability of data and materials

As explained in *Methods*, data generated in *GeoGebra* will not be useful without providing instructions. The links to simulations in *GeoGebra* presented in this study are available with individual instructions from the corresponding author upon reasonable request.

## Funding

There is no funding for this work to report.

## Conflict of interest

There is no conflict of interest related to this work.

## Notes

### Competing Interest Statement

The authors have declared no competing interest.

### Summary of Updates

It updated the section on the configuration space, the original contribution to Listing's law.

